# Why 627 BindCraft2 trajectories produced zero binders

**DOI:** 10.64898/2026.10.13.758722

**Authors:** Wei Fu, Jing Chen, Xu Wu

## Abstract

De novo protein-binder design with deep-learning pipelines such as BindCraft2 has been reported to deliver experimental success rates of 10-100% against well-structured targets. Here we report the opposite tail: two BindCraft2 (v1.0.3) cyclic-peptide campaigns targeting the shallow K81 lactylation epitope of human lactate dehydrogenase A (LDHA) produced zero accepted designs from 627 trajectories. Retrospective analysis of the campaign artifacts shows that 97% of trajectories were rejected at the first (“screen”) stage by a single confidence gate (binder pLDDT < 0.60), not because the interface failed to form: the interface metric i_pTM reached 0.69 and the target was confidently folded (pTM = 0.90). Critically, the mid-project decision to disable the “lysine trick” (forced_targeting) was backwards — that feature moved 17 trajectories past the screen versus 1 without it, and raised interface confidence. We document two preset/metric mismatches that set a macrocycle campaign up to fail regardless of budget: a screen-stage pLDDT gate calibrated for globular binders, and a binder_contacts loss that is degenerate for short peptides. We describe the exact artifacts BindCraft2 writes and distill them into a reusable post-mortem protocol, and we include a complete how-to-use-BindCraft2 guide so the failure analysis also serves as a tutorial.

## 1. Introduction

### 1.1 Protein binder design and BindCraft2

BindCraft2 is an open-source, campaign-based suite for one-shot de novo protein and peptide binder design [1,8]. Its precursor BindCraft was shown to generate nanomolar-affinity binders without high-throughput screening, reporting experimental success rates of 10–100% across receptors, allergens, and CRISPR enzymes [1]. The pipeline interleaves two neural networks inside a differentiable optimization loop: AlphaFold2 (multimer) predicts the structure of the target–binder complex and supplies gradients that steer the binder onto a user-specified epitope [2,3], while ProteinMPNN redesigns the binder sequence to match the evolving backbone [4]. The method belongs to a family of structure-prediction-driven design tools that also includes RFdiffusion [5]. In parallel, deep learning has advanced the design and prediction of short peptides directly — self-assembling-peptide discovery [9], short-peptide aggregation rules [10], and computational peptide and miniprotein inhibitor design [11,12].

### 1.2 The biological target: LDHA K81 lactylation

Lysine lactylation (Kla) is a recently discovered post-translational modification linked to glycolytic flux [6]. The client brief concerns K81 of human LDHA (UniProt P00338; PDB 1I10 [7]), a surface-exposed residue ~26 Å from the NADH nicotinamide. The working hypothesis is that K81 lactylation acts allosterically through the active-site loop (residues 96–107) rather than by direct steric occlusion. The deliverable was 10 cyclic-peptide candidates (head-to-tail macrocycles, 7–16 residues) engaging the K81 neighborhood (residues ~80–85). This is a hard target by construction: the epitope is small and shallow, and a 7–16 residue macrocycle has almost no intrinsic fold — it becomes structured only by engaging the surface.

### 1.3 Scope

This note (i) documents the failed campaigns quantitatively, (ii) explains the root cause from BindCraft2’s own outputs, (iii) corrects a misdiagnosis made mid-project, and (iv) provides a reusable diagnostic protocol and a practical BindCraft2 usage guide. All data and code are listed under Data availability.

## 2. Methods

### 2.1 Target preparation

The design target is chain A of the tetrameric human LDHA structure (PDB 1I10-derived, UniProt numbering, residues 2–332) [7]. The NADH cofactor and pyruvate were stripped so that the surface-epitope design has no buried hetero atoms. Because BindCraft2’s AlphaFold2 representation (atom37) supports only the 20 standard amino acids (plus MSE→MET), the K81-lactyl residue cannot be represented: residue 81 is presented as its parent LYS. The lactyl group occupies a shallow pocket lined by residues 80–85 and E55 (lactyl hydroxyl H-bonds the E55 carboxylate at 3.5 Å); peptides that accommodate the lactyl are therefore identified downstream by explicit MD rather than by the design tool.

### 2.2 BindCraft2 installation

BindCraft2 v1.0.3 was installed at a shared location from the official repository [8]. ‘bash install.sh’ builds a Python 3.12 virtual environment and fetches the AlphaFold2 parameter set and the ProteinMPNN checkpoints. Every session that runs the CLI sources ‘bc2_env.sh’. The preflight step (see 3.9) prints the resolved model-weight paths and is the fastest way to confirm a correct install.

### 2.3 Campaign configuration

Two campaigns were run against the same target, differing only in the forced_targeting property. Run 1 used forced_targeting: true (229 trajectories, aborted manually at 17 h); Run 2 used forced_targeting: false (398 trajectories, four SLURM attempts to completion). Both used the cyclic_peptide modality, binder lengths 7-16, hotspots 80, 82, 83, 84, 85, min_hotspot_contact_final 0.5, and 10 requested final designs. The full JSON is reproduced in Section 3.3. Both campaigns produced zero accepted designs.

### 2.4 Run management

Campaigns were launched via a SLURM wrapper that runs ‘bindcraft design design.json’ under a 23.5 h ‘timeout’ (so the 24 h walltime never SIGKILLs the process) and resubmits itself on a non-zero exit, up to 12 attempts. BindCraft2 resumes a campaign from its ‘.campaign_state.json’ when re-run with the same JSON, so each attempt continues from the accumulated results rather than restarting.

### 2.5 Post-mortem analysis

Three artifact classes were parsed: (i) ‘1_Trajectories/!_Trajectories.csv’ for the terminal stage of each trajectory; (ii) the per-trajectory ‘*_losses.csv’ files for per-round metric curves; and (iii) the worker / design logs for per-trajectory rejection records (stage, reason, and the numeric pLDDT and i_pTM values that triggered the gate). Data were aggregated with Python (csv + numpy). Density estimates use a Gaussian kernel with Silverman’s bandwidth. Figures use the Okabe–Ito colorblind-safe palette; run 1 is blue, run 2 is vermilion.

## 3. Using BindCraft2: a practical guide

This section is a self-contained reference. It reflects BindCraft2 v1.0.3 as observed in this project.

### 3.1 Core concepts

| Term | Meaning |
| --- | --- |
| target | the protein (and chain) the binder must bind; supplied as a PDB |
| binder | the designed molecule; a protein, peptide, or macrocycle |
| modality | a preset of losses and filters: binder, peptide, cyclic_peptide, large_binder, VHH, scFv, Fab, ARP, homo_oligomer, multidomain |
| hotspots | target residues (comma-separated) the binder must engage |
| binder_lengths | [min, max] residue range of the binder |
| trajectory | one full design attempt: fold → redesign → validate |
| campaign | a batch of trajectories under one JSON, accumulated across runs |

### 3.2 The design loop and its six stages

A trajectory passes six ordered stages; each ends in an acceptance gate. The default number of optimization rounds per stage is shown in parentheses.

| Stage | Rounds | Role |
| --- | --- | --- |
| screen | 50 | cheap early fold + first sequence redesign; the first filter |
| refine | 25 | polish the binder fold and interface |
| anneal | 45 | relax the sequence/structure toward the final conformation |
| harden | 5 | confirm the design against stricter confidence gates |
| mutate | 15 | final sequence search |
| final | — | independent validation; writes accepted designs to 3_Ranked/!_Ranked.csv |

### 3.3 A minimal cyclic-peptide design file

The exact JSON used (run 2). Project_folder names the output directory; point it at a new folder to start a fresh campaign.

| Key | Value | Note |
| --- | --- | --- |
| modality | cyclic_peptide | head-to-tail macrocycle preset |
| binder_lengths | [7, 16] | residue range |
| hotspots | 80,82,83,84,85 | K81 neighborhood (K81 excluded) |
| forced_targeting | false | the “lysine trick” (see 3.6) |
| min_hotspot_contact_final | 0.5 | re-adds the $\geq 50\%$ hotspot floor |
| number_of_final_designs | 10 | contract deliverable |
| max_trajectories | 400 | hard budget |
| campaign_seed | 0 | reproducible random draws |
| aa_bias | {"C": 0} | exclude cysteine |
| project_folder | results/kla_epitope | output location |

### 3.4 Losses and weights

The objective is a weighted sum of losses. Defaults in core/default.json (weights_* entries) include the following. Each loss has a documented gradient term and, where relevant, an acceptance filter.

| Loss | Default weight | What it optimizes |
| --- | --- | --- |
| interface_contacts | 1.0 | binder–target contacts (the interface) |
| binder_contacts | 1.0 | long-range intra-binder contacts (see 5.2) |
| binder_pae | 0.4 | intra-binder predicted alignment error |
| interface_pae | 0.1 | binder–target PAE |
| plddt_loss | 0.1 | binder confidence (1 – pLDDT) |
| iptm_loss | 0.05 | interface pTM |
| binder_helicity | –0.3 | negative weight → encourage helicity |
| compactness | 0.5 (0.0 in cyclic preset) | binder compactness |

### 3.5 Filters and gates

Stage gates (min_plddt_* / min_iptm_*) apply during the corresponding stage; the final filters apply to the independently validated design. Values below are the defaults; the cyclic_peptide preset overrides only the final Unbound_Binder_pLDDT and pTM to null (see 5.1).

| Gate / filter | Default | Stage |
| --- | --- | --- |
| `min_plddt_screen` | 0.60 | screen |
| `min_plddt_refine` | 0.60 | refine |
| `min_plddt_anneal` | 0.65 | anneal |
| `min_plddt_harden` | 0.65 | harden |
| `min_plddt_final` | 0.70 | final |
| `min_iptm_anneal / harden / mutate` | 0.50 | anneal / harden / mutate |
| `min_iptm_final` | 0.70 | final |
| `max_ipae_final` | 0.33 | final |
| `Interface_Residues` | $\geq 7$ | final |
| `Backbone_Clashes` | $\leq 0$ | final |
| `Hotspot_Contact_Fraction` | $\geq 0.5$ (if set) | final |

### 3.6 Properties

Properties are optional booleans that add a biological objective plus its filter. The two most relevant here:

- **forced_targeting —** during early gradient stages, every exposed target residue outside a shell (default 10 Å) around the hotspots is mutated to lysine, making the rest of the surface unattractive so the binder is steered onto the epitope; the true sequence is restored before validation. It steers where the binder lands, not how tightly it binds.
- **disulfide_staple —** adds a geometric disulfide filter and allows cysteine; an alternative cyclization chemistry to head-to-tail closure, useful when a macrocycle fails to close or fold.

Other properties include humanize, protease_stable, termini_accessible/together, mixed_topology, and the objectives induced_fit, fold_switch, and multidomain.

### 3.7 Autotuner and desperation ladder

Two mechanisms adapt a stalled campaign automatically. The autotuner (on by default) nudges the screen and refine stage lengths every ten trajectories. The desperation ladder (on by default) triggers after ‘desperation_trajectories’ (default 750) trajectories with nothing accepted, then trades away difficulty one rung at a time (initial_guess → target_flexibility → held-out multimer validation → raised recycles). Each rung raises the expected false-positive rate, so ladder-accepted designs should be weighted accordingly.

### 3.8 Output files

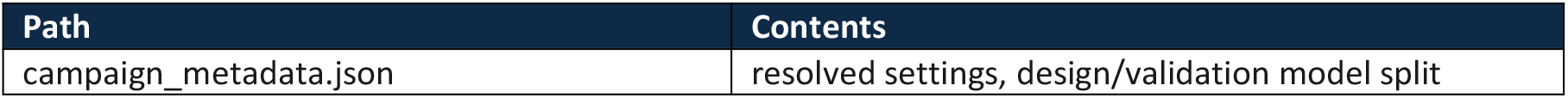
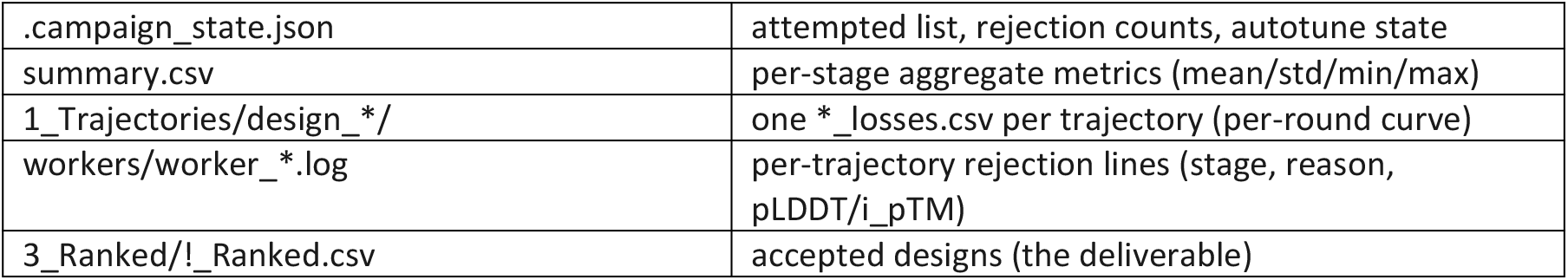

### 3.9 Pitfalls we hit

- Re-running the same JSON resumes, not restarts — use a fresh project_folder for a new campaign.
- Long campaigns outlive any walltime; wrap the CLI in timeout and resubmit on non-zero exit.
- Validate first: a preflight run on a login node (no GPU) catches bad settings before burning queue time.
- Short-peptide modalities inherit protein-scale stage gates — check, and relax, the stage pLDDT gates (see 5.1).

## 4. Results

### 4.1 The stage funnel

Of 627 trajectories, 609 (97.1%) terminated at the screen stage — the first gate. Only 18 reached any later stage, and none was accepted.

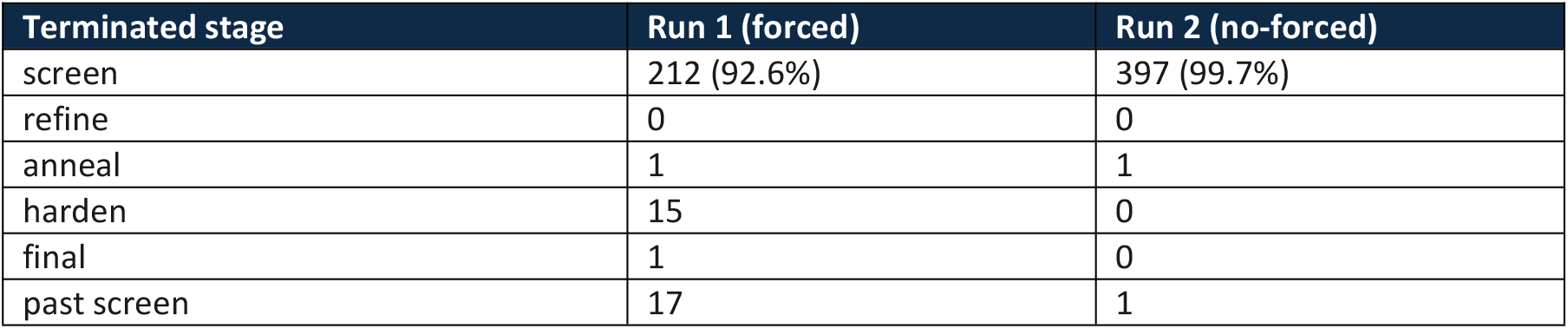

**Figure 1.**
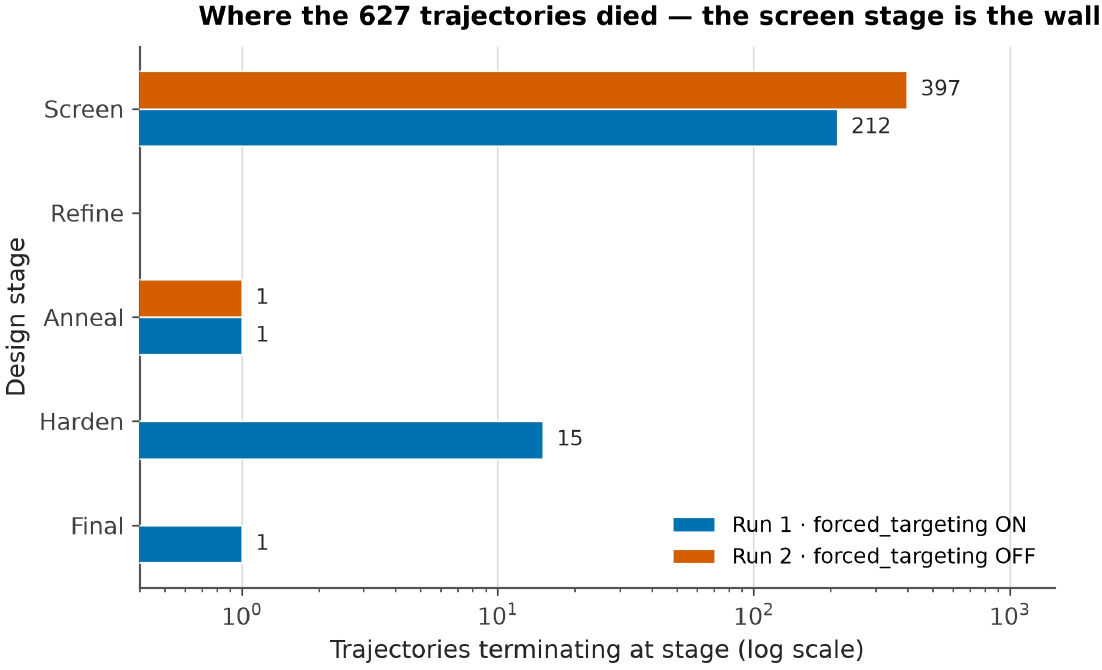
Stage funnel. Bars are trajectories terminating at each design stage (log x-axis). Nearly all trajectories die at the first (screen) gate.

### 4.2 Confidence at rejection

The screen stage compares the binder’s predicted pLDDT against min_plddt_screen = 0.60. The distribution of that metric at the moment of rejection never clears the gate.

| Metric (at rejection) | Run 1 (forced) | Run 2 (no-forced) |
| --- | --- | --- |
| binder pLDDT | 0.24–0.64 (mean 0.38) | 0.28–0.63 (mean 0.35) |
| interface i_pTM | 0.10–0.69 (mean 0.32) | 0.16–0.65 (mean 0.25) |
| rejection reason | pLDDT (213); i_pTM+pLDDT (15) | pLDDT (398) |

**Figure 2.**
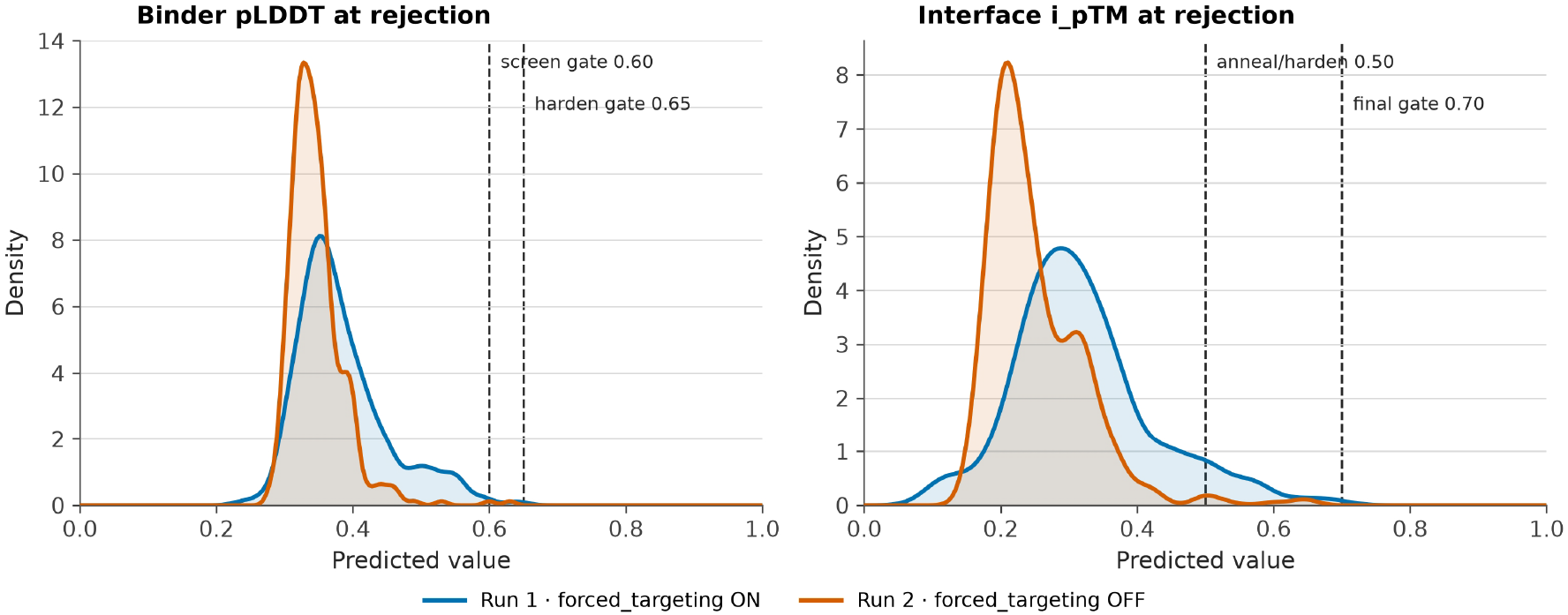
Density of the two confidence metrics at the point of rejection (Gaussian KDE, Silverman’s bandwidth). Left: binder pLDDT sits below the 0.60 screen gate. Right: interface i_pTM rarely approaches the 0.70 final gate. Dashed lines mark the gates.

### 4.3 The empty acceptance corner

Plotting each trajectory in (pLDDT, i_pTM) space shows the acceptance region (pLDDT ≥ 0.60 and i_pTM ≥ 0.70) is empty. The two runs separate along the interface axis: forced targeting shifts points toward higher i_pTM, while both runs remain pinned below the pLDDT wall.

### 4.4 Target confidence

The target itself was confidently folded throughout: pTM was 0.90 (run 2) and 0.75 (run 1; the lower value is a design-time artifact of forced_targeting, which temporarily mutates the target surface to lysine). The binder made a handful of interface contacts (mean ≈ 4.3–4.9 residues within cutoff) but never attained a confidently predicted fold. In short, the protein was fine and the peptide was floppy, and AlphaFold said so loudly.

## 5. Discussion

### 5.1 The screen pLDDT gate is miscalibrated for macrocycles

min_plddt_screen = 0.60 is inherited from the generic core profile and tuned for globular binders. A 7–16 residue cyclic peptide rarely has a self-confident fold in AlphaFold’s eyes — its structure is largely induced by the target. Applying a protein-scale confidence gate at the very first stage guarantees that macrocycle designs are culled before the interface objectives can act. Notably, the cyclic_peptide preset already relaxes the final Unbound_Binder_pLDDT and pTM filters to null for exactly this reason, but leaves the stage gates untouched — an inconsistency that, in effect, was the whole campaign.

**Figure 3.**
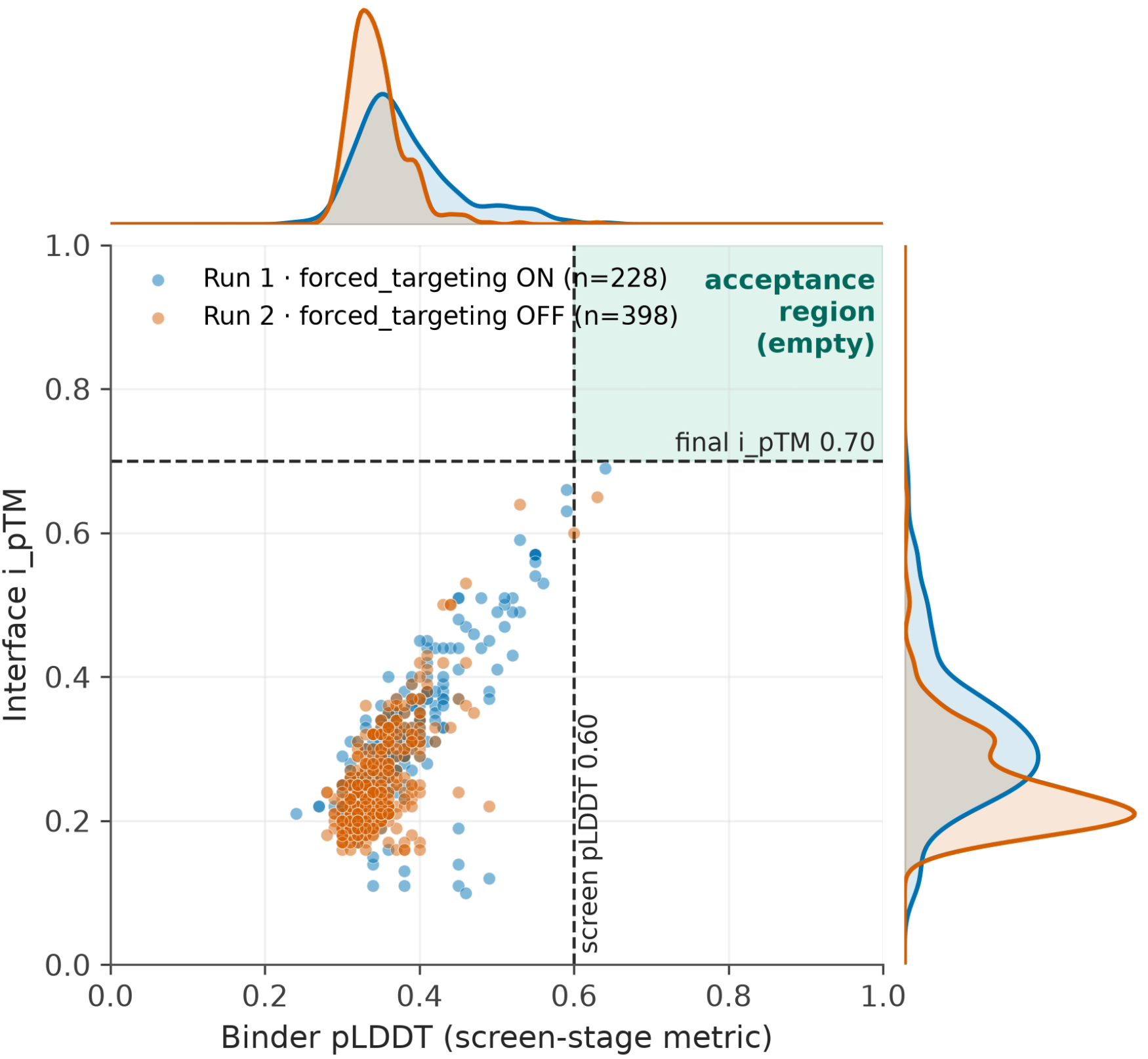
Joint plot: scatter of pLDDT vs i_pTM with marginal densities. The green shaded region is the acceptance corner — empty. Run 1 (blue) reaches higher i_pTM than run 2 (vermilion) but neither crosses the 0.60 pLDDT gate.

### 5.2 A metric that says nothing: binder_contacts

Every one of the 627 trajectories recorded binder_contacts = 0.0. This is not evidence of an unfolded peptide; it is evidence of a degenerate loss. The default binder_contacts loss only scores residue pairs separated by ≥ 9 positions in sequence. For a 7–16 residue binder that leaves almost no evaluable pairs, so the loss is trivially satisfied (or empty) for every trajectory. Lesson: for short-peptide modalities, binder_contacts is a silent no-op — do not read “zero contacts” into a zero value, and do not expect it to drive folding.

### 5.3 forced_targeting helped — the mid-project diagnosis was backwards

Run 1 (forced_targeting on) was aborted on the hypothesis that the “lysine trick” was poisoning the designs. The artifacts disagree. Run 1 moved 17 trajectories past the screen (including one to the final stage) versus 1 for run 2; its interface confidence was higher (i_pTM mean 0.32 vs 0.25); and its rejection reasons are richer — 15 trajectories died at harden on [i_pTM, pLDDT] after passing screen, whereas run 2’s 398 rejections are all [pLDDT] at screen. Forced targeting was doing its job — steering the binder onto the epitope — and its only cost (a lower target pTM) is restored before validation. The correct lever was the confidence gate, not the targeting.

### 5.4 Is the epitope undesignable as posed?

The residual signal is real and worth owning: even the best i_pTM (≈ 0.69) falls below the 0.70 final floor. A shallow, solvent-exposed pocket of five residues offers a short macrocycle little to wrap around, and with no induced structure there is no confident fold to predict. This reflects a general property of short peptides — weak intrinsic folding and hard-to-predict aggregation behavior [10] — which is why computational peptide and miniprotein inhibitor design remains an active field [11,12]. The actionable conclusion is not that the pipeline is broken, but that this target needs a different pose — a longer binder, a stapled/disulfide-cyclized scaffold, or a broader hotspot set. The pipeline, run correctly, tells you which of those to try next (Section 6).

### 5.5 Implications for BindCraft2 users

- For short peptides, treat the stage pLDDT gates as a tuning parameter, not a fixed default.
- Build the stage funnel first — it localizes the failure to a single gate before any settings change.
- Change one variable between campaigns and keep results in separate folders for a clean comparison.

## 6. A reusable post-mortem protocol

The entire Section 4 was reconstructed from three artifacts BindCraft2 always writes. The recipe:

1. Build the funnel — read 1_Trajectories/!_Trajectories.csv (the terminated column) and count trajectories per stage. If > 90% die at one stage, that stage’s filter is the bottleneck.
2. Read the rejection reasons — parse the worker/design logs for “rejected at <stage> … due to [<filters>]”. The bracket names the exact gate(s) that fired.
3. Plot the rejection metrics — the same log lines carry the numeric values. A distribution pinned below a gate is a calibration problem (relax the gate); one that straddles the gate is a genuine design problem (change epitope/length).
4. Check for degenerate metrics — per-trajectory losses that are constant (our binder_contacts = 0.0) are not signals.
5. Change one variable between campaigns, keep the seed/format comparable, and keep results separate.
6. Only then touch the scientific knobs — binder_lengths, hotspots, cyclization route, or initial_guess — using the stage funnel as your before/after metric.

## 7. Conclusion

A failed campaign, if its artifacts are read correctly, is more informative than a silent success. Here the artifacts show a campaign that was never given a chance: a confidence gate calibrated for globular proteins culled 97% of short-peptide trajectories before the interface could mature, and a misdiagnosis sent the second run in the wrong direction. We are now testing a third campaign with macrocycle-appropriate stage gates and initial_guess enabled; if it too returns zero, the data has already told us the next move — the epitope itself, not the pipeline, is the variable to change.

